# Campaign-Based Citizen Science for Environmental Mycology: the “Science Solstice” and “Summer Soil-stice” Projects to Assess Drug Resistance in Air and Soilborne *Aspergillus fumigatus*

**DOI:** 10.1101/2020.06.11.146241

**Authors:** Jennifer M. G. Shelton, Matthew C. Fisher, Andrew C. Singer

## Abstract

Citizen science projects are often undertaken for ecological and environmental research purposes but also have great potential for use in microbiology research to track the emergence and spread of pathogens in the environment. ‘Science Solstice’ and ‘Summer Soil-stice’ are mycology citizen science projects aimed at assessing drug resistance in *Aspergillus fumigatus* fungal spores found in air and soil, respectively, in the United Kingdom (UK). *A. fumigatus* plays an important role in the environment as a decomposer of plant material, but is also an opportunistic human lung pathogen. Infection with drug-resistant spores can lead to a worse clinical outcome for the patient.

On the first four solstice and equinox days between June 2018 and June 2019, volunteers were asked to collect air samples from their homes and workplaces and return them to our lab in Freepost envelopes. An additional round of samples was requested from volunteer’s gardens and/or compost on the June 2019 solstice. In total, 787 volunteers returned 2,132 air samples and 509 soil samples, which grew a total of 7,991 *A. fumigatus* colonies. The estimated total cost of the study was £2,650; the equivalent of 33 pence per *A. fumigatus* colony grown.

Incorporating citizen science into the environmental surveillance of drug-resistant *A. fumigatus* allowed for the simultaneous collection of hundreds of environmental samples across the entire UK on the same day. The insights generated from this study would not be practical in the absence of public participation and offers opportunities to ask scientific questions that were previously unaskable.

## Introduction

Citizen science is defined as the “intentional involvement, in a non-professional capacity, of people in the scientific process, e.g. the collection … of data” (Pocock, 2015) and is becoming increasingly popular for simultaneously conducting research and engaging with the public about science. Many citizen science projects in the UK rely on volunteers to monitor population levels of native insects (Gardiner, 2012; Lye, 2012; Wilson, 2018), wildlife (Hof and Bright, 2016), birds (Cannon, 2005; Sparks, 2017) and plants (Rich and Woodruff, 1990; Pescott, 2015). Citizen scientists can also report environmental incidents with potentially harmfully effects such as toxic algal blooms (Ransom Hardison, 2019) or river pollution (Hyder, 2017), and can aid surveillance of invasive species (Pocock and Evans, 2014), wildlife diseases (Robinson, 2010; Lawson, 2012) or plant pathogens (Brown, 2017). The majority of these projects ask participants to record their observations, either online, through an app or via post over a prolonged period of time.

Projects may also raise awareness of invisible health threats such as air pollution, pathogen spread and antimicrobial resistance (AMR). One example is Netherlands-based iSPEX, where participants measured atmospheric aerosols on a single day using an optical add-on for their smartphones with a corresponding app that collated data (Snik, 2014). A UK-based example is Swab & Send: an ongoing, self-funding microbiology project asking citizen scientists to take swabs of any object or environment they choose to help identify new antibiotic compounds (www.lstmed.ac.uk/public-engagement/swab-send). Public health-focused projects like these provided the inspiration for this study. We asked volunteers to collect samples from the air and soil, at home and work in the UK, for the surveillance of antifungal-resistant spores of *Aspergillus fumigatus*, a ubiquitous decomposer of dead plant matter and opportunistic human lung pathogen.

On average, we inhale 100s of *A. fumigatus* spores a day (Kwon-Chung and Sugui, 2013), some of which cause hypersensitisation and “fungal asthma” or aspergillosis disease ranging from chronic colonisation of the airways to invasive bloodstream infections. In the UK, as many as 400,000 individuals suffer from severe asthma with fungal sensitisation (SAFS), approximately 238,000 individuals with aspergillosis lung disease and an estimated 4,200 individuals with invasive aspergillosis (IA) (Pegorie, 2017). IA has a mortality rate ranging from 30-80% (Bongomin, 2017) and its prevalence is increasing in the UK due to increasing numbers of patients receiving immunosuppressive therapies for transplant, cancer or autoimmune conditions and the ageing population (Löbermann, 2012). Patients that are in critical care with severe viral infections such as influenza are at high risk of IA (Schauwvlieghe, 2018), and we are already witnessing examples of IA in patients that are ill with COVID-19. Increasingly, these infections are resistant to the medical antifungals (i.e. azole drugs) used to treat them despite no prior exposure of the patient to these drugs, suggesting environmental acquisition of resistance by the infecting spores pre-inhalation. Early diagnosis and treatment are associated with better patient outcomes, yet a survey by The Aspergillosis Trust revealed that diagnosis took between 1 to 5 years for 60% of the 128 respondents (personal comms, Sandra Hicks and Gillian Fairweather at The Aspergillosis Trust). In a survey of the scientific community on Twitter (*n* = 1,267; April 2020), only 54% replied “yes” when asked if they had heard of aspergillosis or knew what it was. This study aimed to raise awareness amongst participants by publishing blog posts on institute websites and including information sheets in sampling packs about *A. fumigatus*, aspergillosis and the relevance of widespread environmental sampling.

To date, much of the focus around environmental monitoring of airborne *A. fumigatus* spores in the UK has been on industrial composting facilities and potential risks to workers and nearby residents, with reports published by Department for Environment, Food & Rural Affairs (Defra) (Knight, 2009), Environment Agency (EA) (Environment Agency, 2018) and Health & Safety Executive (HSE) (Gilbert, 2003). Further studies have collected air and/or soil samples from areas in the UK over time to assess the prevalence of azole-resistant *A. fumigatus*: Greater Manchester from 2009-2011 (Alshareef and Robson, 2014; Bromley, 2014), Dublin from 2014-2016 (Dunne, 2017), South Wales from June to November 2015 (Tsitsopoulou, 2018) and 6 sites across Southern England from May to July 2018 (Sewell, 2019). These studies give valuable insight but are limited in sample number and coverage due to sample collection being undertaken by the study authors themselves. In order to address some of these problems, this study reports the UK-wide collection of outdoor air and soil samples by citizen scientists, in a campaign-based, single timepoint manner, from which *A. fumigatus* spores were cultured and will ultimately be tested for azole antifungal-resistance.

## Methods

The aims of this citizen science project were to monitor for drug-resistant *A. fumigatus* spores in outdoor air and soil across the UK at multiple timepoints. The ‘citizen science’ methodology of the study had several rationales: 1) to provide a step-change in UK spatial coverage from previous studies, 2) to raise awareness of aspergillosis diseases amongst the general public, and 3) to trial the efficacy of the chosen sample collection methods on citizen scientists as a viable approach for mycological research. The outcome of the study will be to determine whether there are spatial or temporal determinants of resistance that could inform future policies to protect those at risk for aspergillosis.

We thereby asked individuals residing in the UK to collect spore samples from their local air on four dates (21^st^ June 2018, 24^th^ September 2018, Friday 21^st^ December and 20^th^ March 2019) and garden soil on one date (21^st^ June 2019). These dates were chosen because they: 1) were the solstice and equinox dates making them easy to remember for participants and ‘catchy’ for the purpose of advertising; 2) were equally spaced throughout the year making it useful for examining seasonal shifts in spore recovery; and 3) allowed for sufficient time in the laboratory to process samples before the next sampling campaign.

### Recruitment for citizen science projects

Participants were recruited for the projects by posts published on social media platforms Twitter and Facebook, as well as on several mycology websites and The Aspergillosis Trust website (www.aspergillosistrust.org), containing a poster (Figure 1), a brief description of the project and a link to a Google form (Supplementary Figure 1). Printed posters were displayed outside author and co-author offices and on noticeboards around Imperial College London (ICL) and UK Centre for Ecology & Hydrology (UKCEH). The posters displayed the name of each project, all of which incorporated solstice or equinox, and an image of Stonehenge, which is iconic for such celestial events; all in an effort to make the sampling dates memorable. The posters also contained a brief description of the project that aimed to be understandable to non-scientists, links to an online blog post containing further information about the project, and the Twitter handles of the author and project to be followed for regular updates. Twitter was chosen as a way of providing project updates because Tweets are visible to the public and did not require the participants to befriend or follow the authors as on other social media platforms. Twitter updates also avoided potentially upsetting participants by sending unsolicited emails. At the bottom of each poster was a shortened URL to the Google form. Emails were sent by co-authors to ICL and UKCEH mailing lists containing a description of the project and a link to the Google form.

**Figure 1:**
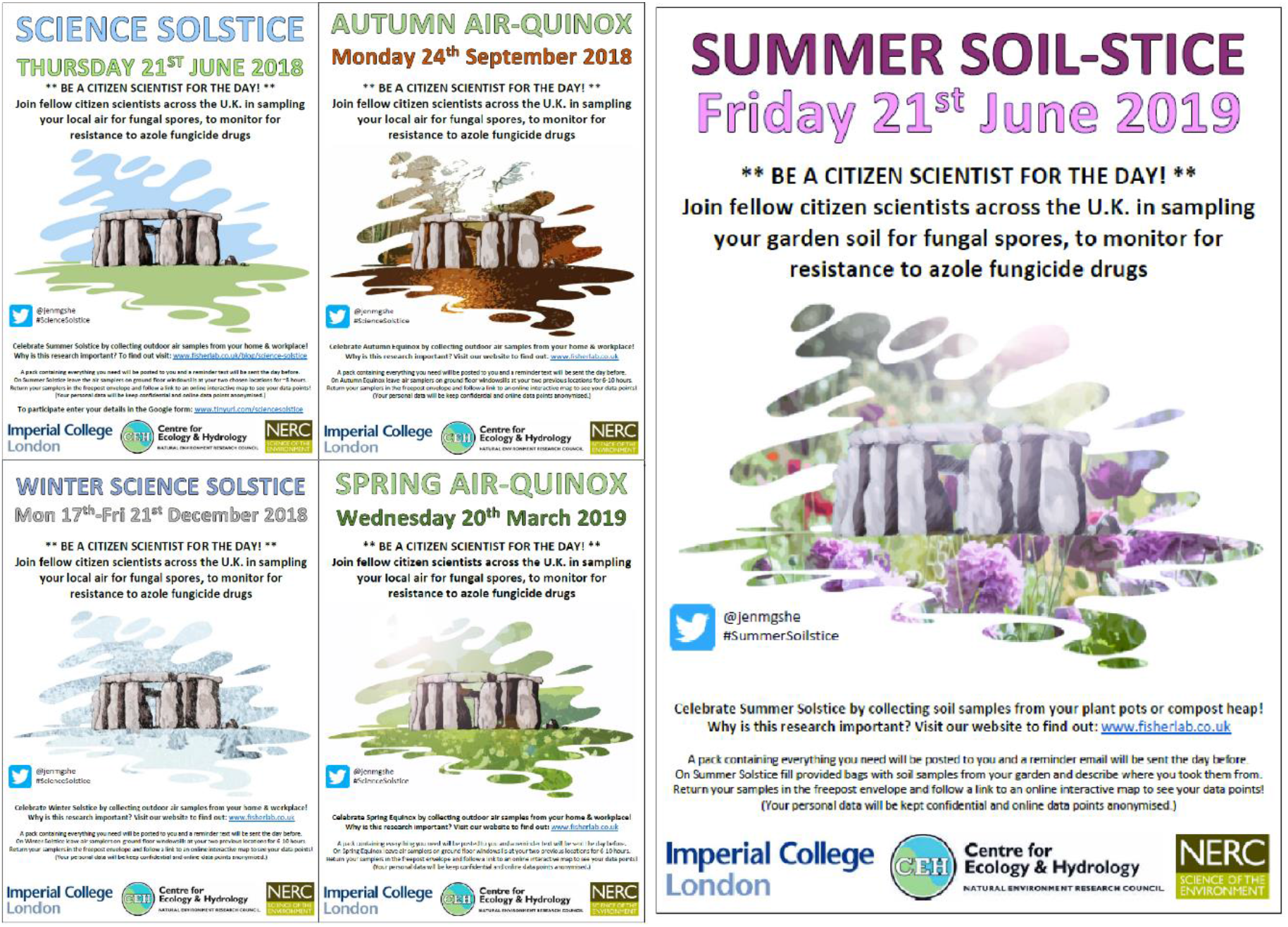
Posters advertising for citizen scientists to take part in UK-wide air and soil sampling projects were displayed around ICL and UKCEH and posted on social media platforms prior to sampling dates.

On the Google form, participants were requested to provide their name and address, for postage purposes, their email address for a reminder email sent the week before and mobile number for a reminder text to be sent the evening before. For the initial air sampling round, optional questions asked them for the research institute they are affiliated to (if any) and to say how they had heard of the project. All communications were checked for General Data Protection Regulation (GDPR)-compliance as of new rules introduced on 25^th^ May 2018 and participants were informed *via* the Google form and *via* email about how their personal data would be used, stored and kept confidential.

### Air sampling citizen science

Participants who filled in the Google form to take part in one or more of the four air sampling rounds were sent an air sampling pack containing a MicroAmp™ clear adhesive film (Applied Biosystems™, UK) cut in half (to produce two “air samplers”) with adhesive putty attached for securing in place. The pack also contained a poster (Figure 1), a questionnaire (Supplementary Figure 2), simple instructions (Supplementary Figure 3) and a brief scientific description of the experimental aims (Supplementary Figure 4). Participants were asked to attach the air samplers to outdoor ground floor windowsills at their home and workplace and expose by peeling off the backing slip for 6-10 hours on sampling day. If torrential rain was forecast, participants were asked to sample on the soonest dry day after as rain falling on the air samplers reduced their stickiness and therefore their ability to capture spores. They were then asked to re-cover the air samplers and return them by post, along with the completed questionnaire, in the Freepost envelope provided. The questionnaire asked the date and geographical locations of sample collection, whether they were collected from outdoor ground-floor windowsills, and for the participant to provide an email if they wished to receive updates. Upon receipt, *A. fumigatus* colonies were cultured directly from the air samplers onto petri dishes containing agar and stored at 4°C in a refrigerator for further analysis.

### Soil sampling citizen science

Participants who filled in the Google form to take part in soil sampling, which followed after the air sampling rounds, were sent a pack containing two plastic sachets, a wooden spatula, a poster (Figure 1), simple instructions (Supplementary Figure 5) and a brief scientific description of the experimental aims (Supplementary Figure 4). They were asked to fill two plastic sachets with soil from their garden and complete a questionnaire (Supplementary Figure 6) detailing the geographical location of their garden, the location of the soils within their garden (pot or planter, border, bag of compost, bag of manure, compost heap) and a brief description of the sample (e.g. plant or bulb type in pot, brand of compost or manure, contents of compost heap). They were then asked to return the sealed sachets of soil and the questionnaire in the Freepost envelope provided. Upon receipt, 2 g of each soil sample was plated onto petri dishes containing agar to culture *A. fumigatus* colonies, which were then stored at 4°C in a refrigerator for further analysis, along with the remainder of the soils.

For the soil sampling project, the blog post published on the UKCEH website (Supplementary Figure 7) explained that compost heaps and bags of compost might act as “hotspots” for the growth of azole-resistant *A. fumigatus*. In an effort to mitigate exposure, participants were advised to exercise caution when sampling from these locations as disturbance can lead to aerosolization of large numbers of spores. People were asked not to take part in the project if they suffer from aspergillosis, have a lung condition (chronic or acute, such as ‘flu) or are immunosuppressed, as these all put them at greater risk of contracting aspergillosis from inhaling a large number of spores. Participants were asked to sample from locations within their own garden only so they experienced equivalent or lesser exposures in taking part as from standard gardening activities such as potting, digging and compost manipulation. Participants were free to opt out at any time by emailing the primary author or by not collecting samples.

### Citizen science engagement

Participants were encouraged throughout the projects to share photos of their sampling on the designated day via Twitter or email. When provided, participants were asked for their consent for this material to be used in future work and presentations by the author. Participants were also given the option to opt-out of the projects at any time by emailing the author or by not returning their samples. Participants were asked on the questionnaires to indicate whether they were happy to receive future project updates by email. Those who opted to receive updates were sent an email approximately 4-6 weeks after each round when all samples had been processed thanking them for their participation, informing them of the number of samples received, the number of *A. fumigatus* colonies grown and a link to an online Google map showing the location of each sample processed and the number of colonies grown from it.

On 25^th^ June 2018, four days after the initial air sampling round, an email was sent to participants who provided an email address asking for feedback on the project via a different Google form (Supplementary Figure 8). This form asked the reason participants did not take part (if they didn’t), whether the email and text reminders were useful, whether they’d like to part again if the experiment was repeated, and had a comment box for additional feedback.

## Results

### Citizen science participation in the UK

Across the four air sampling projects spanning June 2018 to March 2019 a total of 485 unique individuals residing in the UK collected one or more air samples. A total of 1,293 air sampling packs were sent out and 976 were returned, equating to an overall participation rate of 75%. Participants collectively returned 1,896 air samples across the four dates, which were collected from all over England, Northern Ireland, Wales and Scotland. Screenshots of the Google maps sent out to participants after the air and soil sampling rounds is shown alongside a UK population density map in Figure 2, which shows that the majority of samples were sent in from populous areas. Concomitantly, areas with the lowest coverage of air and soil sampling are also less densely populated. Results for individual air sampling rounds and for the soil sampling round are shown in Table 1.

**Figure 2:**
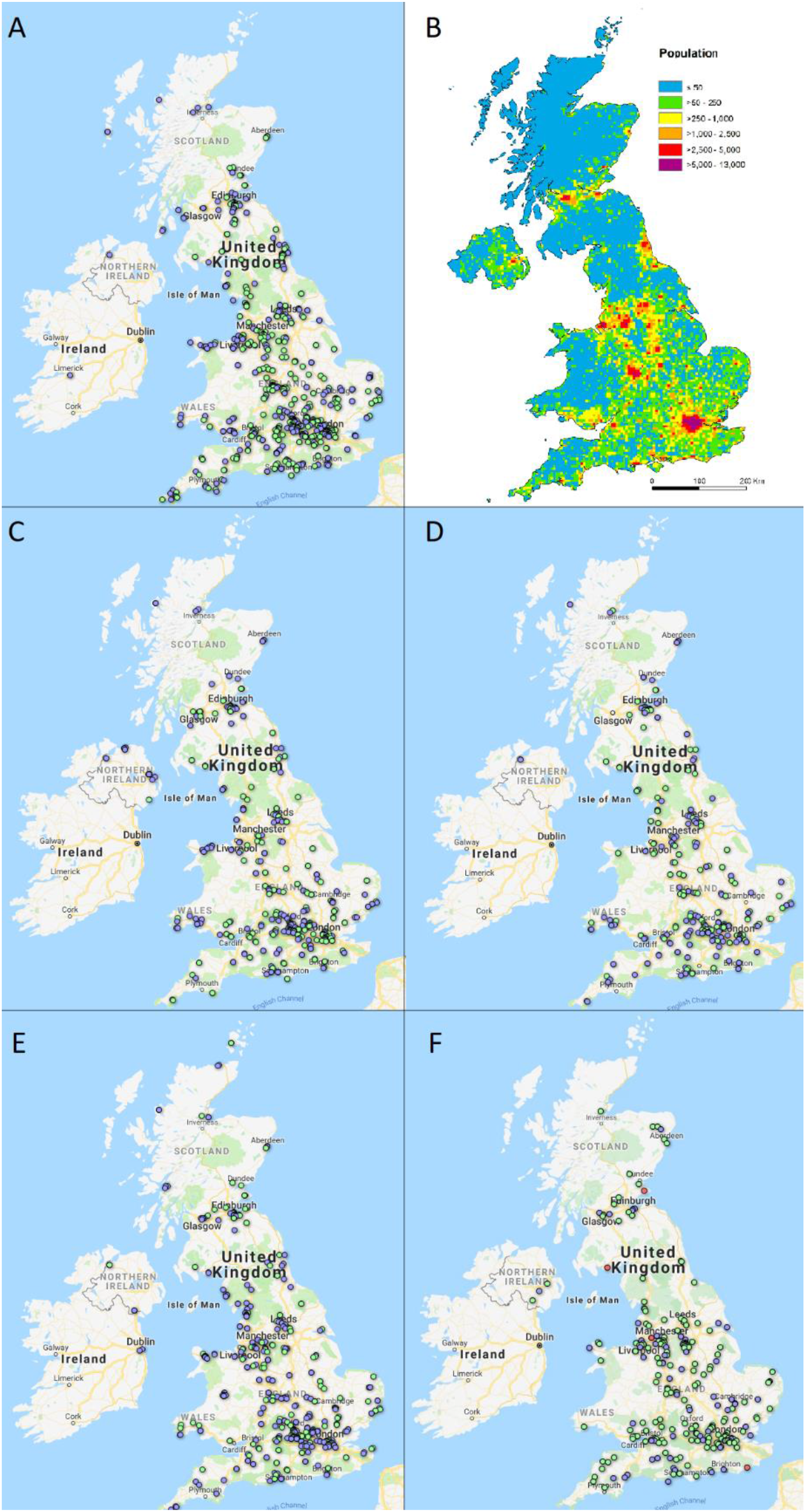
Google maps showing locations that participants collected air samples from on A) 21^st^ June 2018, C) 24^th^ September 2018, D) 21^st^ December 2018 and E) 20^th^ March 2019. F) shows locations that soil samples were collected from on 21^st^ June 2019. Blue dots indicate samplers that did not grow *A. fumigatus* colonies, green dots indicate samplers that did and red dots indicate samplers that were contaminated with other fungal growth. B) is a population density map of the UK produced by Vieno et al (2015) (Heal and Williams, 2015).

**Table 1:** Numbers of sampling packs sent out and number of packs and samples returned across the four air sampling dates and one soil sampling date.

| Type of sampling | Sampling date | Packs sent out | Packs returned | Return rate (%) | Samples collected |
| --- | --- | --- | --- | --- | --- |
| Air | 21<sup>st</sup> June 2018 | 461 | 365 | 79 | 712 |
|  | 24<sup>th</sup> September 2018 | 300 | 204 | 68 | 398 |
|  | 21<sup>st</sup> December 2018 | 231 | 165 | 71 | 321 |
|  | 20<sup>th</sup> March 2019 | 301 | 242 | 80 | 465 |
| Soil | 21<sup>st</sup> June 2019 | 334 | 246 | 74 | 509 |

Of the 365 participants in the first air sampling round, 160 (43%) also took part in the second round, 120 (32%) in the third round and 112 (30%) took part in all four air sampling rounds. Of the 246 participants in the soil sampling round, 43 (17%) had already taken part in one or more of the air sampling rounds.

### Citizen science participation globally

Due to the global nature of Twitter the first air sampling round attracted 52 participants from 16 countries in addition to the UK. Whilst global sampling was not the intention of this study, sampling packs were sent to these participants for comparative analysis to UK samples. For the first air sampling round global participants sent back a total of 144 samples from Australia, Belgium, Canada, Chile, China, France, Germany, Hungary, Italy, Madagascar, New Zealand, Portugal, Spain, The Gambia, The Netherlands and USA. The second air sampling round received 92 samples from 50 individuals overseas: Canada, France, Germany, New Zealand, Portugal, Spain and USA. For the third and fourth air sampling rounds it was decided not to send sampling packs abroad as the Freepost return envelopes were not valid in other countries and the authors thought it unfair for participants to pay for postage. Soil sampling was open to UK participants only due to restrictions on moving soil samples between countries.

### Isolation of *A. fumigatus* from samples

The 1,896 air samples collected and returned from the UK across the four air sampling rounds grew a total of 2,366 of fungal colonies that were identified morphologically as *A. fumigatus*, and the 236 air samples collected globally across the first and second air sampling rounds grew a total of 451 *A. fumigatus* colonies (Table 2). The 509 soil samples grew a total of 5,174 colonies. The average number of colonies per air sample ranged from 1.8 to 3.1 across the four sampling rounds, whereas the average number per soil sample was 15.8.

**Table 2:** Numbers of samples that grew *A. fumigatus* colonies and the number of colonies cultured across the four air sampling rounds and one soil sampling round. ^a^ These numbers represent fungal isolates cultured from samples that morphologically resembled *A. fumigatus*.

| Type of sampling | Sampling date | Number of samples that grew *A. fumigatus* (% of total)<sup>a</sup> | Number of *A. fumigatus* colonies cultured<sup>a</sup> | Average number of colonies grown from *A. fumigatus*-positive samples |
| --- | --- | --- | --- | --- |
| Air | 21<sup>st</sup> June 2018 |  |  |  |
|  | UK | 408 (57) | 1152 | 2.8 |
|  | global | 81 (56) | 280 | 1.9 |
|  | 24<sup>th</sup> September 2018 |  |  |  |
|  | UK | 190 (48) | 429 | 2.3 |
|  | global | 63 (69) | 171 | 1.9 |
|  | 21<sup>st</sup> December 2018 | 152 (47) | 477 | 3.1 |
|  | 20<sup>th</sup> March 2019 | 169 (36) | 308 | 1.8 |
| Soil | 21<sup>st</sup> June 2019 | 327 (64) | 5174 | 15.8 |

### Recruitment method

For the initial air sampling project participants were asked several optional questions on the Google sign up form. Of the 513 individuals who completed the form, in the UK and globally, 233 (45%) belonged to a research institute and 489 (95%) indicated how they’d heard of the project. The research institutes that recruited the most individuals were the author’s institutes UKCEH (*n* = 36) and ICL (*n* = 16). The ways that individuals heard of the project were on: Facebook (*n* = 139), email (*n* = 128), Twitter (*n* = 103), word-of-mouth (*n* = 97) and other (*n* = 46). The Tweet recruiting individuals for the first air sampling round (#ScienceSolstice) made 27,731 impressions and received 642 total engagements. The Tweet recruiting for subsequent air sampling rounds (#AutumnAirquinox, #WinterScienceSolstice and #SpringAirquinox) made 13,288 impressions and had 244 engagements, and the Tweet for soil sampling (#SummerSoilstice) made 29,350 impressions and had 823 engagements.

### Adherence to sampling date amongst UK samples

The initial air sampling round had the highest adherence to sampling date of 94%, which dropped to ~60% for the second and fourth air sampling round (Table 3). This drop was due to the author’s communication with participants preceding the second, third and fourth air sampling dates to collect air samples on days either side of the sampling date if they were unable to participate on the sampling date itself. The decision was taken that sample number was more important than sampling date, as feedback from the first air sampling round suggested that flexibility in sampling date would increase participation. The third air sampling round had the lowest adherence to date because it was the week before Christmas and the weather was unpredictable so participants were encouraged to sample on any date between 17^th^ and 21^st^ December 2018 with suitable weather, which 274 (85%) did. Sample date was less important for the soil sampling project because it was not affected by weather conditions, so less emphasis was placed on timing for this sampling round.

**Table 3:** Dates that UK samples were collected for four air sampling rounds and one soil sampling round. ^a^ Due to weather conditions, and proximity to Christmas, participants for the third air sampling round were asked to collect on any suitable day between 17^th^ – 21^st^ December, hence low adherence to sampling date.

| Type of sampling | Sampling date | Samples missing date (% of total) | Number of samples collected on intended sampling date (% of total) | Range of sampling dates |
| --- | --- | --- | --- | --- |
| Air | 21<sup>st</sup> June 2018 | 8 (1) | 669 (94) | 20<sup>th</sup> June – 1<sup>st</sup> July |
|  | 24<sup>th</sup> September 2018 | 9 (2) | 249 (62) | 17<sup>th</sup> September – 1<sup>st</sup> October |
|  | 21<sup>st</sup> December 2018<sup>a</sup> | 4 (1) | 12 (3) | 12<sup>th</sup> December – 19<sup>th</sup> January |
|  | 20<sup>th</sup> March 2019 | 5 (1) | 287 (61) | 13<sup>th</sup> March – 24<sup>th</sup> April |
| Soil | 21<sup>st</sup> June 2019 | 2 (0) | 261 (51) | 16<sup>th</sup> June – 22<sup>nd</sup> July |

### Participant feedback

During all air and soil sampling rounds, participants engaged with the author by emailing or tweeting photos of themselves or family taking part in the sampling (Figure 3). The author received many messages of support by email, on Twitter and handwritten on completed questionnaires and has been requested by several participants to talk about the citizen science projects in schools and at meetings and conferences.

**Figure 3:**
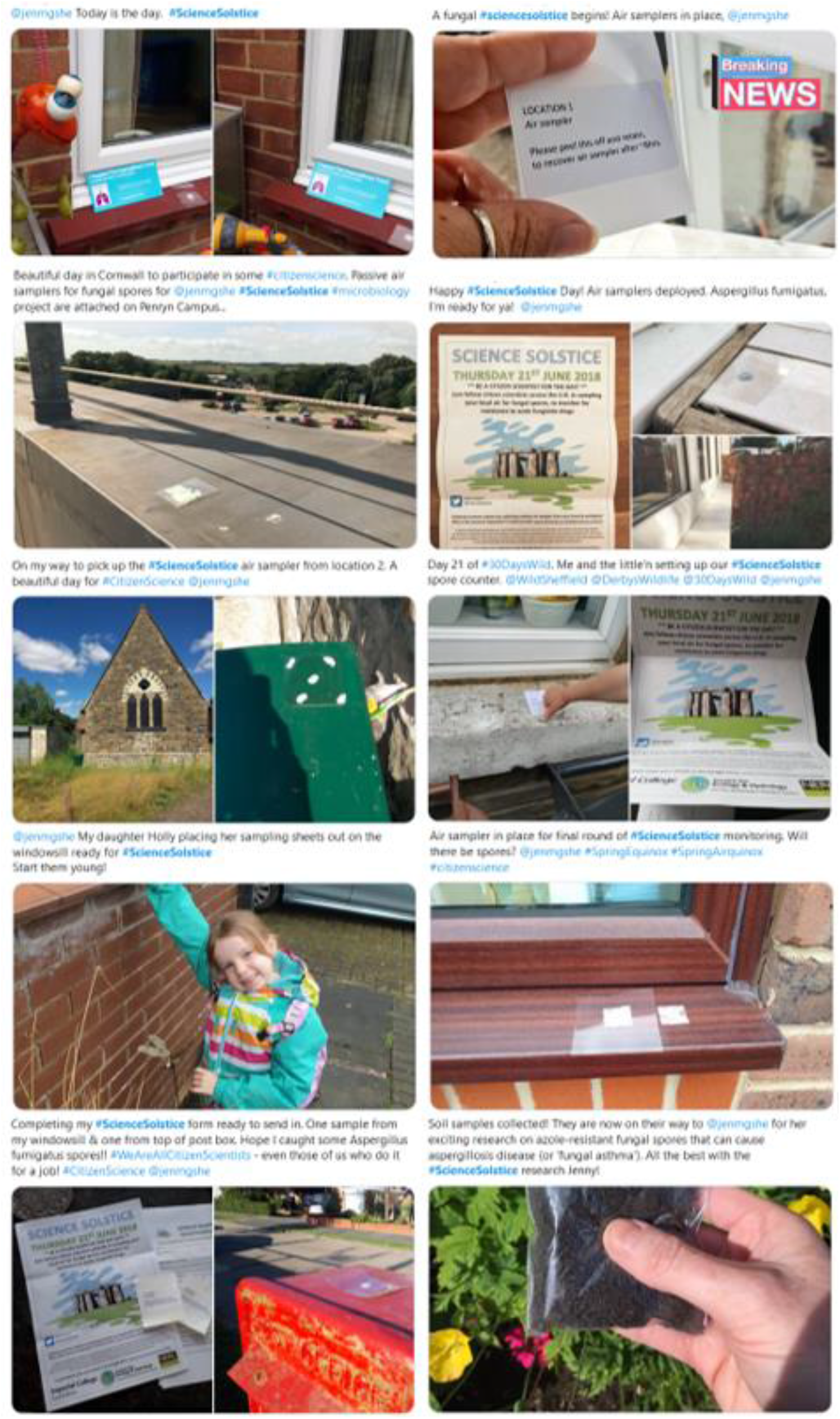
Several Tweets posted by participants on sampling days showing them taking part in air and soil sampling. (Permission granted by participants for these Tweets to be displayed.)

After sending update emails for each sampling round, the primary author received email replies from participants expressing their interest in the results, pleasure in taking part and even disappointment or apologies that their sample(s) had not grown *A. fumigatus*. Leading up to, and following on from, these update emails were progress updates on Twitter that included photographs of samples being returned and the primary author in the laboratory processing samples. These Tweets were well-received and maintained contact with participants who had opted to follow the primary author and/or project on Twitter, without sending unsolicited emails to participants who had only agreed to receive the final update email. They also allowed interested individuals to follow the project even if they hadn’t participated, and on several occasions these individuals got in touch asking to participate in future sampling rounds.

There were 118 responses to the Google feedback form emailed out after the first air sampling round and only 9 responses were from individuals who had not participated, for the following reasons: air sampler(s) blew away or was removed (*n* = 6), air sampling pack did not arrive in time (*n* = 1) and they hadn’t made it to a post box yet but still intended to post (*n* = 2). 115 responders (98%) found the email reminders helpful and 2 (2%) did not receive any. 98 responders (85%) found the text reminder (Supplementary Figure 9) the evening before useful, 3 (3%) did not find it useful, and 14 (12%) did not receive it. 115 of the 118 responders said they would like to participate in the citizen science experiment again if it were repeated and provided their email address. 79 responders left additional feedback in the comment box (Supplementary Table 1), which was overall very positive and encouraging.

Suggestions for improvements were made by participants in personal communications with the primary author and *via* the Google feedback form (Supplementary Table 1) and amendments were made to subsequent sampling rounds. In the first air sampling round white sticky labels that read “LOCATION 1 (or 2) air sampler: please peel this off and retain, to re-cover air sampler after ~8hrs.” were stuck to the back of each air sampler (Figure 4A), to correspond with locations 1 and 2 on the questionnaires. It quickly became apparent on 21^st^ June 2018 that several individuals had removed this label instead of peeling off the backing of the air sampler. These individuals were contacted immediately to remedy this mistake, when possible, but 20 air samplers were returned that had not been exposed correctly. It is possible to tell when an air sample has not been exposed because it appears white and debris-free whereas exposed air samples are yellow, dust and debris is visible and the sticky face and backing slip are misaligned. For subsequent air sampling rounds the sticky labels were not used and air samplers were instead labelled by hand either “LOCATION 1” or “LOCATION 2” (Figure 4B) and participants referred to the instruction page for exposing the air samplers.

**Figure 4:**
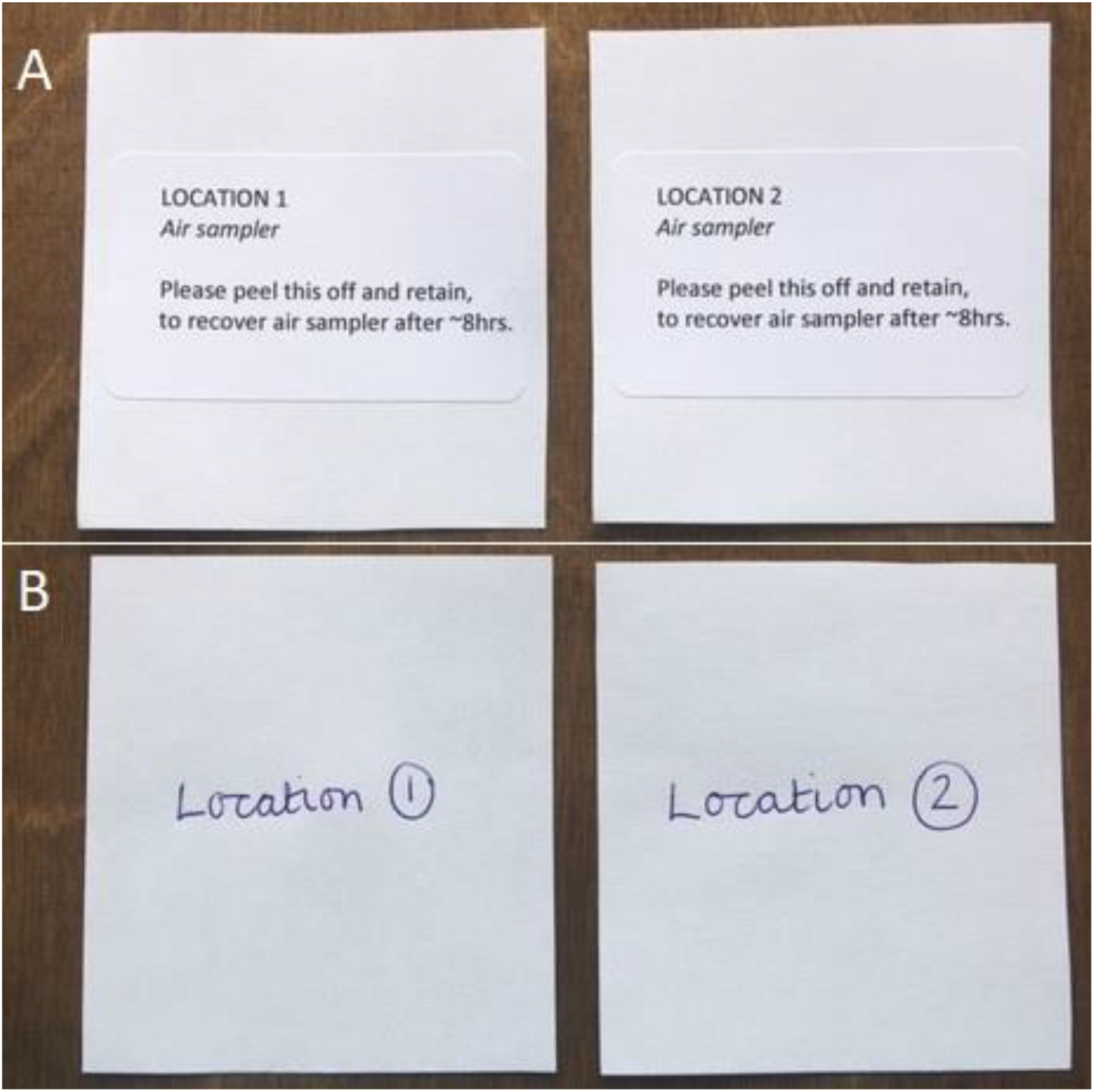
A) Air samplers for first air sampling round had sticky labels on the back with location number and basic instructions, but several participants pealed this off instead of the back of the air sampler. B) For subsequent air sampling rounds location number was handwritten on the back of each air samplers, to avoid confusion on how to expose the air sampler.

Participants also commented during and after the first air sampling round that their air samplers had blown away because adhesive putty was insufficiently adhesive, so for the second air sampling round the author instead attached double-sided foam tape to each air sampler. After the second air sampling round participants reported that the foam tape remained stuck to their windowsills and required chemical removal, so for the third and fourth air sampling round the author provided both adhesive putty and foam tape for participants to choose between.

Several participants contacted the author before, during and after the first air sampling round to apologise for not taking part because they were occupied on the sampling date. As a result, the author amended correspondence for subsequent air sampling rounds asking participants to collect samples on the sampling date whenever possible, but to collect them within 3 days either side of the sampling date if more convenient. A timeline of changes made to sampling methodology as a result of participant feedback is shown in Supplementary Figure 10.

## Discussion

This study asked citizen scientists to collect air and soil samples on five dates between June 2018 and June 2019 and subsequently received a total of 2,132 air samples and 509 soil samples from 787 individuals. Advertising the first air sampling round on social media platforms Facebook and Twitter achieved a high initial enrolment due to the willingness of individuals to share posts, such that the initial Tweet containing the poster and a link to the Google sign-up form reached over 27,000 people. Individuals recruited through Facebook were mostly friends and family of the author, those recruited by email were colleagues at ICL or UKCEH and those who signed up through Twitter were colleagues, fellow mycologists, followers of The Aspergillosis Trust and the general public. Subsequent air sampling rounds benefitted from high retention rates, which participants told the author in personal communications was due to the simplicity of the sampling method, relatively small time commitment required, the enjoyment in taking part and their interest in results.

Organising sampling rounds to take place on single dates, which coincided with solstice and equinox days, was easier for the author and likely achieved a higher return rate than ongoing sampling. The author was able to send an email reminder several days before each sampling date and a text reminder the evening before such that, combined with television and radio broadcasts on solstice and equinox days, no participants stated they’d not taken part due to having forgotten about the project. The single sampling dates also meant that the majority of samples were returned in the following fortnight so the author was able to prepare and use lab consumables within a short timeframe, which is helpful for sterile culturing in mycology. Adherence to sampling date was exceptionally high (94%) for the first air sampling round and moderate for the second and fourth air sampling rounds (~60%) but dropped for the third (12%) due to winter weather conditions and proximity to Christmas. It is worth noting that the weather for the first, second and fourth air sampling rounds (summer, autumn and spring, respectively) was remarkably good, which likely increased participation levels and adherence to sampling dates. Importantly, the range of sampling dates did not overlap between sampling rounds so they can still be ascribed to different seasons as intended. Furthermore, every participant recorded the date of sampling on their questionnaire so, when undertaking spatial analyses in future, the authors can download meteorological data and adjust for factors such as wind speed and wind direction on the day each sample was collected.

The cost of sample collection by citizen scientists was considerably lower than if the author had attempted to collect this number and distribution of samples alone. Recruitment by social media platforms and email was free, stationery was provided by UKCEH and ICL and sampling packs were sent out using ICL’s franking system, so major costs were the setup and renewal of Royal Mail first-class Freepost return and purchase of lab consumables. Additionally, for the first and second air sampling rounds, global participants were reimbursed for postage costs when possible but global participation was discontinued in subsequent sampling rounds. An estimated £1,800 was spent on laboratory reagents and consumables, £50 on stationery and £800 on Royal Mail postage costs bringing the total cost of this project to £2,650. This is the equivalent of £1 per sample or 33 pence per *A. fumigatus* isolate collected across all air and soil sampling rounds, both from the UK and globally.

The majority of participants across the five sampling rounds were resident in the UK (87%) and samples were received from England, Northern Ireland, Scotland and Wales. Without the involvement of citizen scientists, it would not have been possible for the authors to collect this number and distribution of air and soil samples at five single time points. The participation of the author’s friends, family and colleagues show up as clusters of samples around Hampshire/Dorset (author’s home place), Oxfordshire (UKCEH) and London (ICL), which contrasts with under-sampling in central Wales, central Scotland and Northern Ireland. The density of samples overlaps strongly with population density, so these clusters might be due to the under-sampled areas being less densely populated. The sampling discrepancies do not impact the microbiology or genetics aspect of the author’s future work, but might hinder spatial analyses. Spatial coverage in low-population density regions would need to be addressed if the study was repeated.

The authors’ reflection on this citizen science approach for sample collection is that it has exceeded our expectations in terms of participation levels, distribution of samples, timing of sample collection and numbers of *A. fumigatus* colonies grown for onward analysis. Involving citizen scientists has been an incredibly rewarding experience, because of their messages of support, Tweets and photos of them taking part in sampling, invitations to speak at schools and conferences and general enthusiasm. The authors hope this study has also raised awareness of aspergillosis diseases amongst participants both in the UK and globally. This study has resulted in a collection of 7,991 *A. fumigatus* isolates that are to be tested for susceptibility to azole drugs to determine the prevalence and distribution of azole-resistance here in the UK. The sampling techniques used were simple, inexpensive and standardized and could potentially be adapted and used to monitor environmental levels of other fungal, bacterial or viral pathogens, DNA or toxins, insects or chemicals of interest.

## Acknowledgments

The authors would like to thank all the citizen scientists who took part in this study; for contributing samples and for their enthusiasm throughout. We also thank Kathy Richards for her assistance with data entry, Roseanna Collins for helping to process soil samples in the laboratory, and Joshua Jackson and Chris Edwards for their assistance with laboratory administration. Thank you to Michael Pocock for his helpful comments on an earlier version of this manuscript.

## Author Contributions

JMGS, ACS and MCF conceived the study and recruited participants. JMGS processed samples and communicated with participants. JMGS drafted the original manuscript, which ACS and MCF reviewed and provided edits.

