## Supplementary Files for "Campaign-Based Citizen Science for Environmental Mycology: the “Science Solstice” and “Summer Soil-stice” Projects to Assess Drug Resistance in Air and Soilborne *Aspergillus fumigatus*"

Supplementary Figure 1: Google sign-up form for the first air sampling round that was linked to by a shortened URL on the poster and in all recruitment Tweets, Facebook posts and emails.

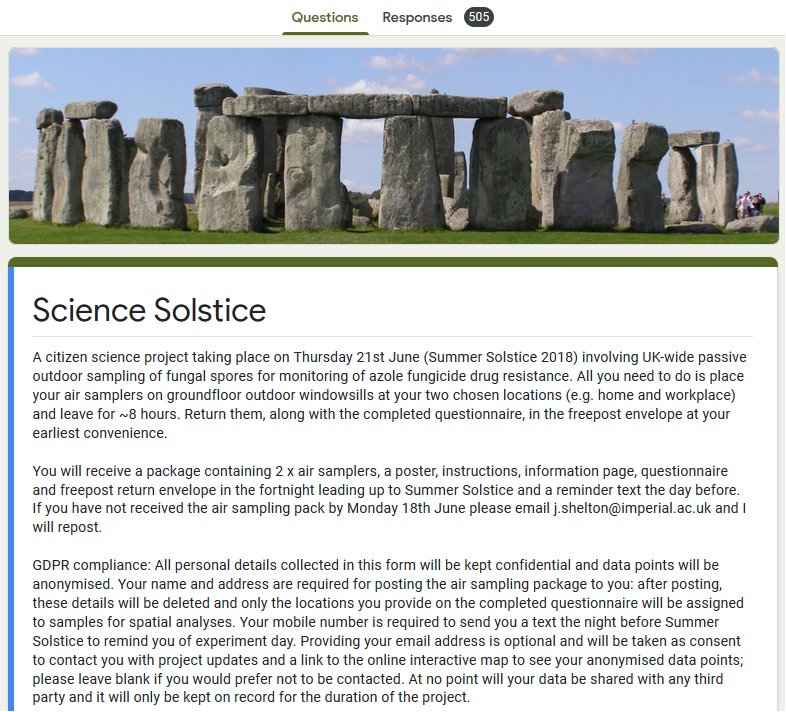

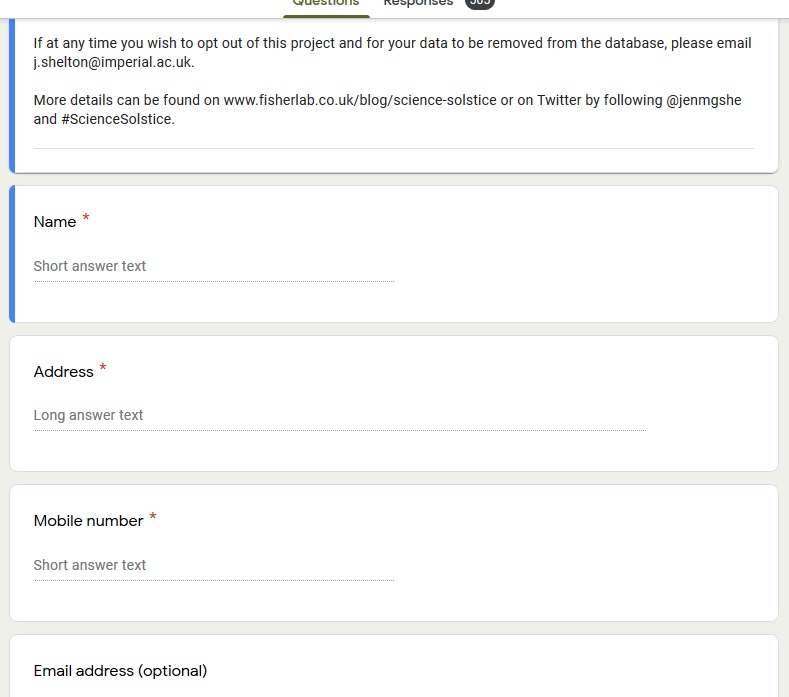

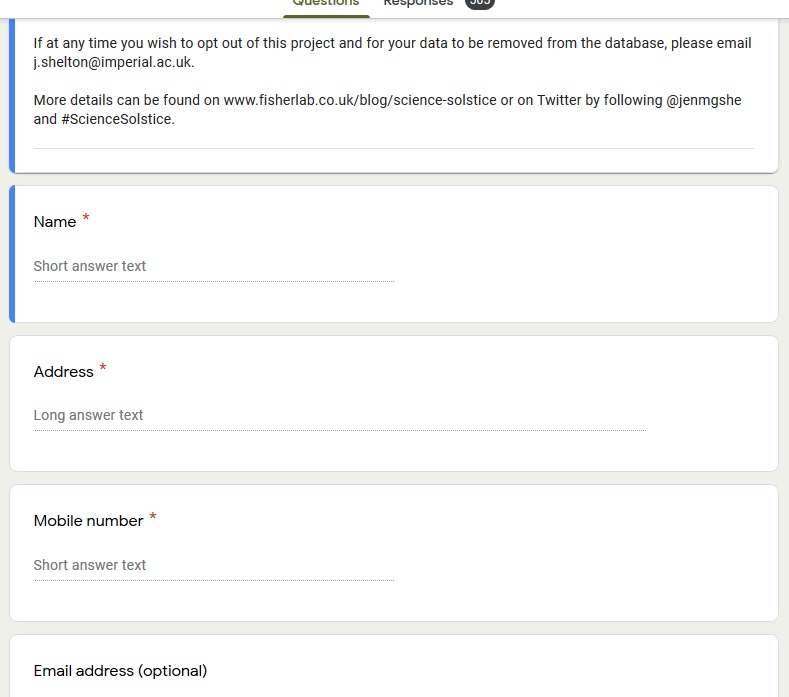

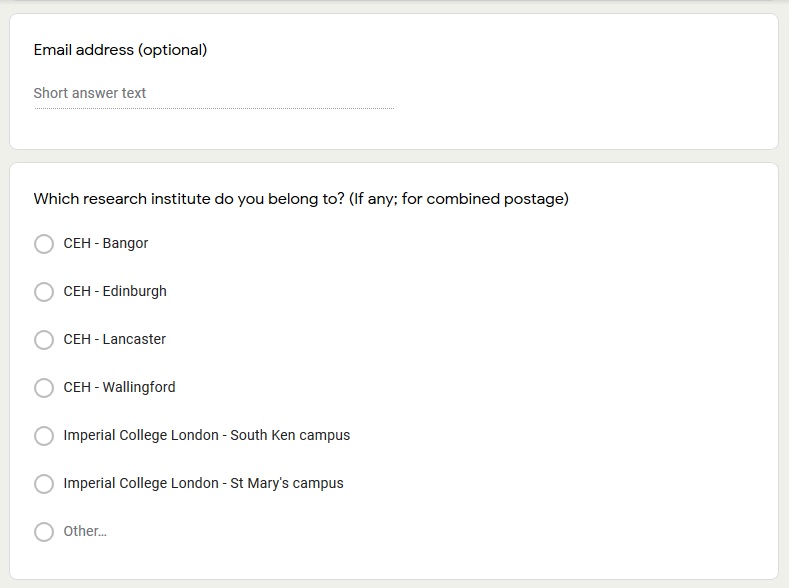

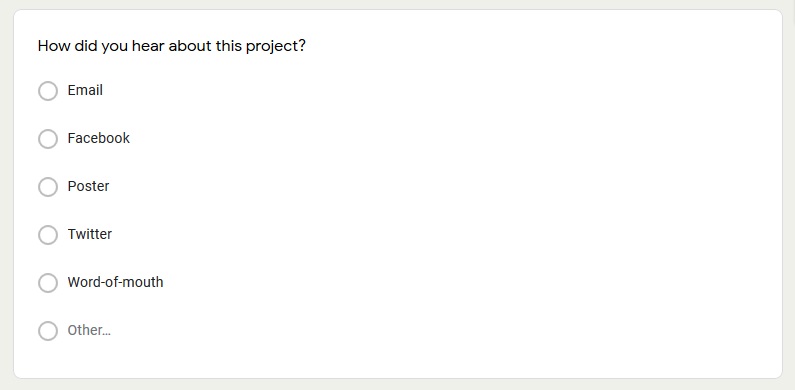

Supplementary Figure 2: Questionnaire to be completed and returned in the freepost envelope detailing the date air samples were collected and the exact locations where the air samplers were placed and exposed. Participants were asked to provide an email address if they wished to received future project updates.

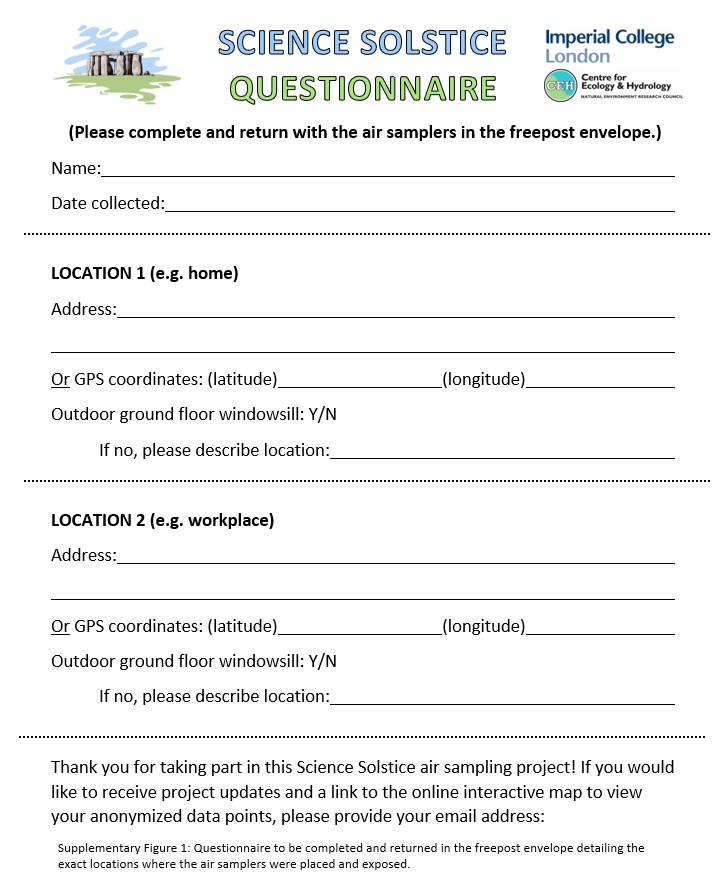

Supplementary Figure 3: Instructions explaining when and how to expose the air samplers to participate in the citizen science experiment. These instructions are for initial air sampling round on 21^st^ June 2018 and the date was amended for subsequent sampling rounds.

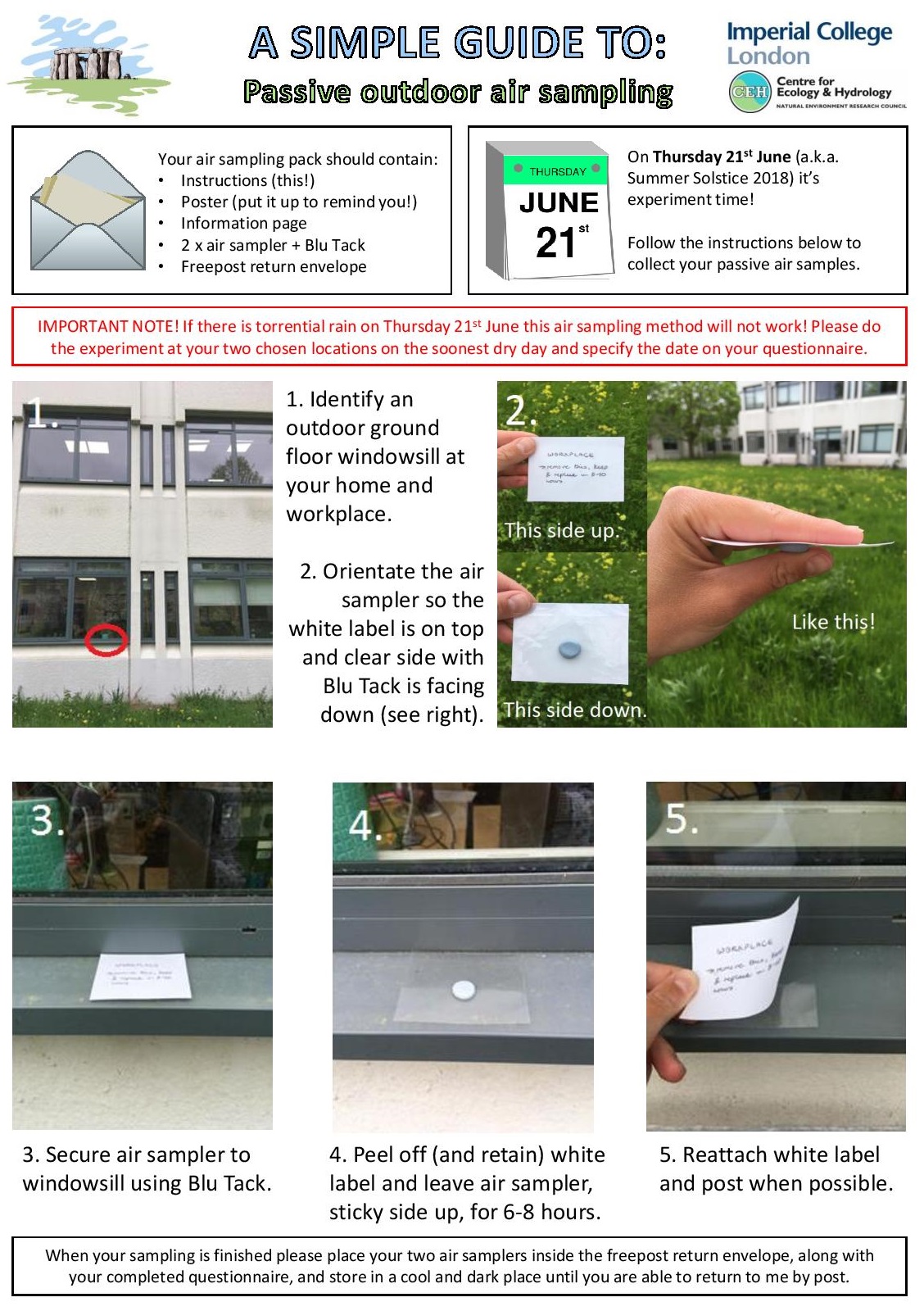

Supplementary Figure 4: A blog post written about the citizen science project was published online when recruitment began, and a paper copy was included in each sampling pack.

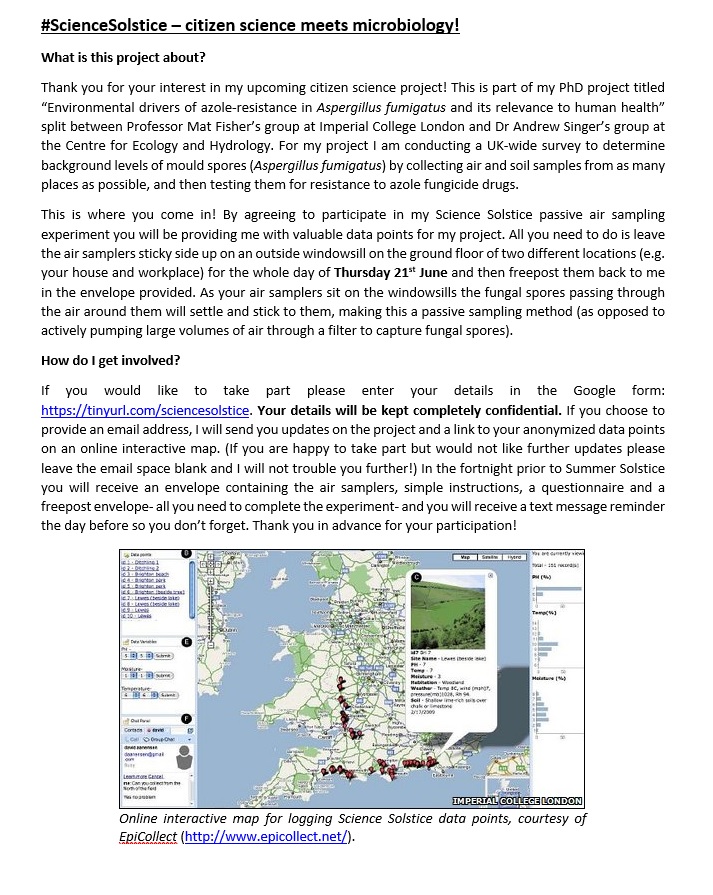

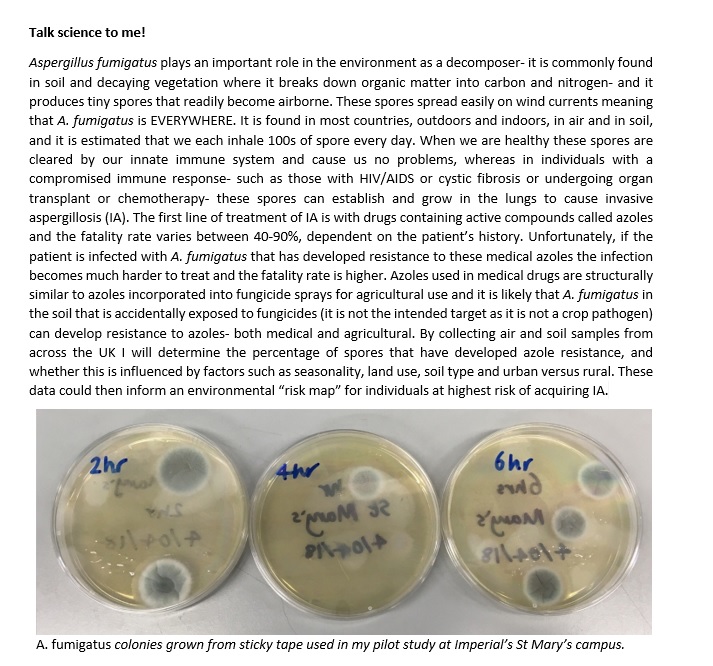

Supplementary Figure 5: Instructions explaining how to collect soil samples to participate in the citizen science experiment.

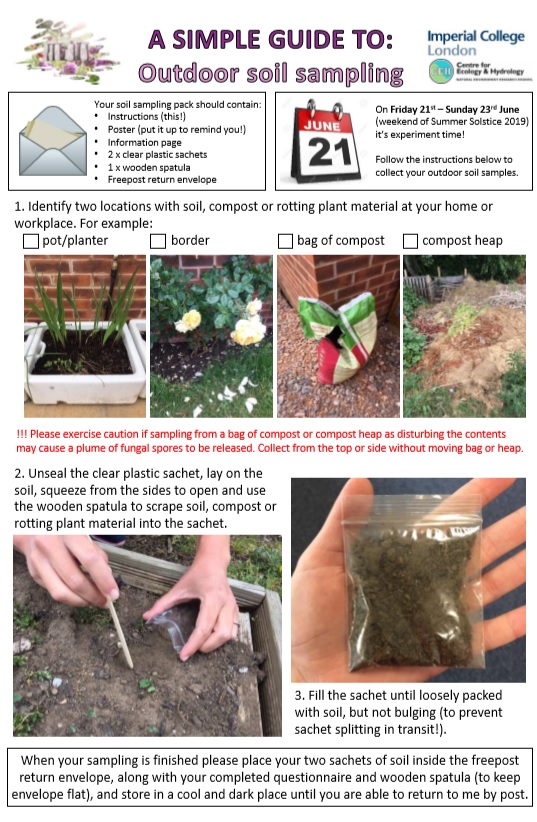

Supplementary Figure 6: Questionnaire to be completed and returned in the freepost envelope detailing the date sample collected, geographical location of the garden that soils were collected from, type of sample collected and sample details.

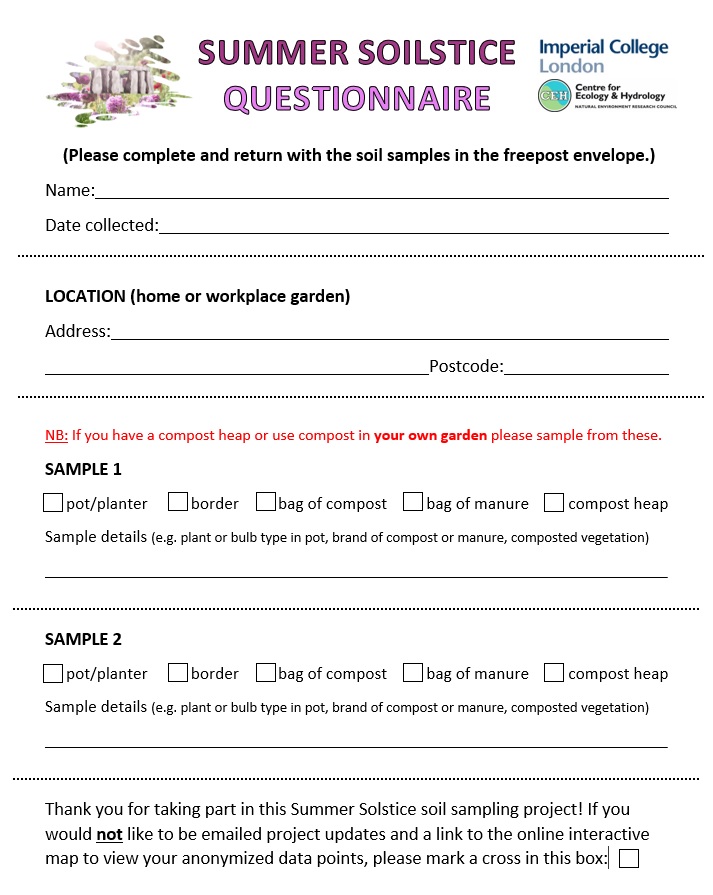

Supplementary Figure 7: Blog post about soil sampling citizen science project published on UK CEH website.

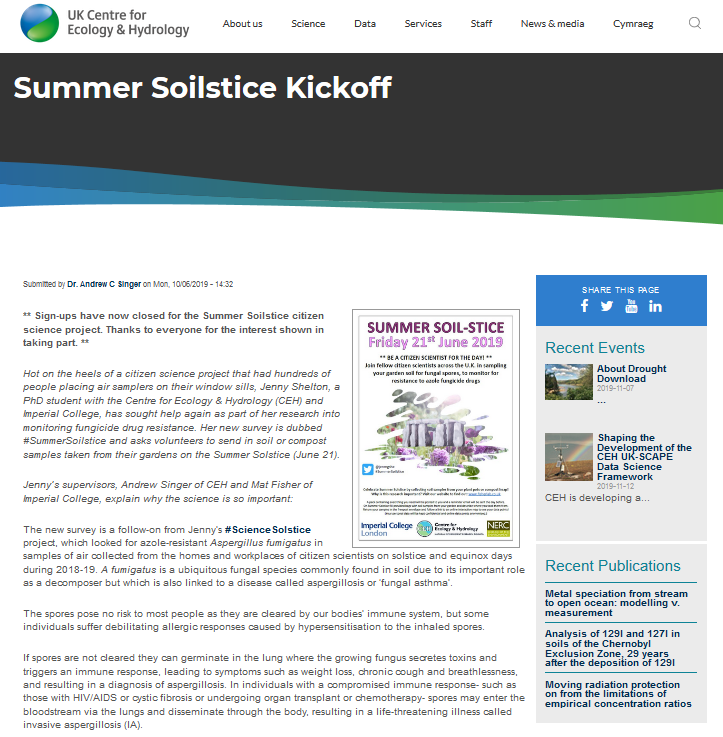

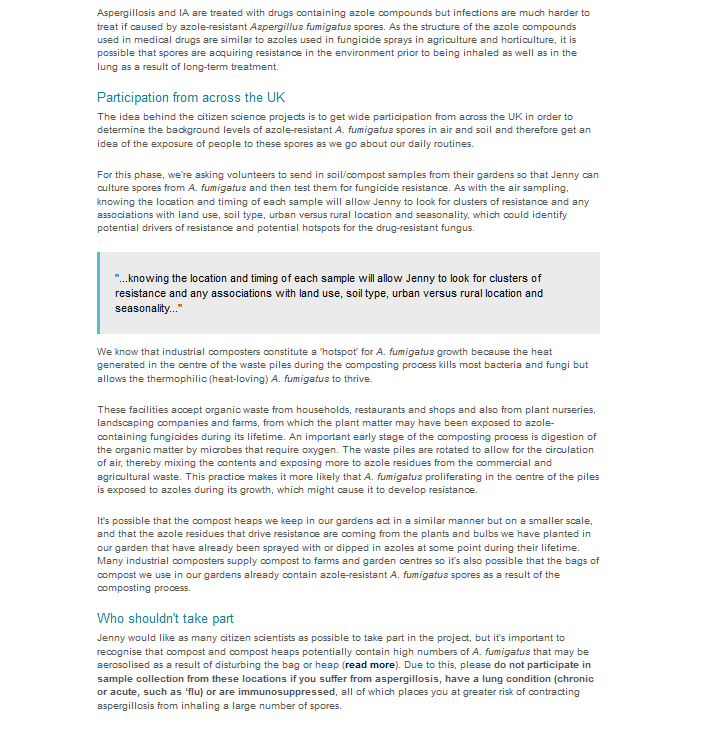

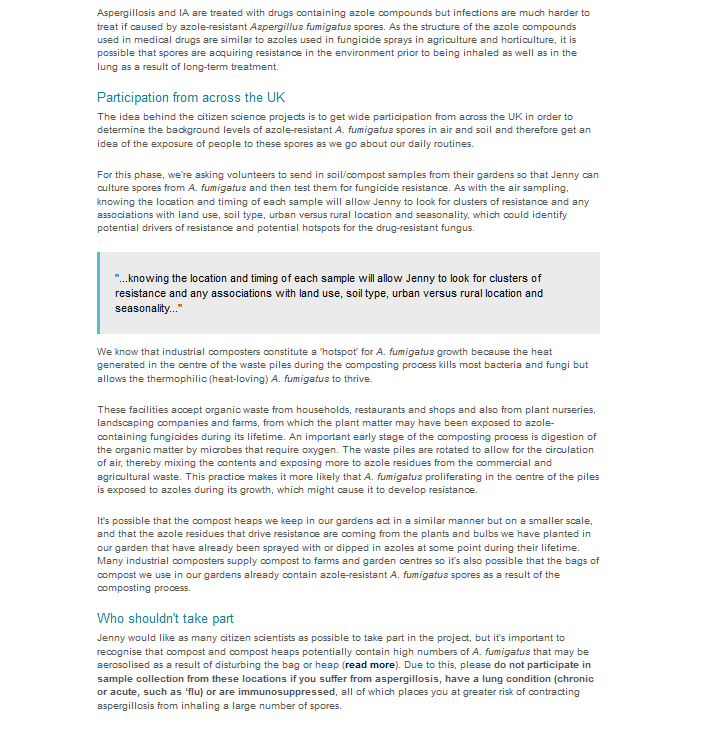

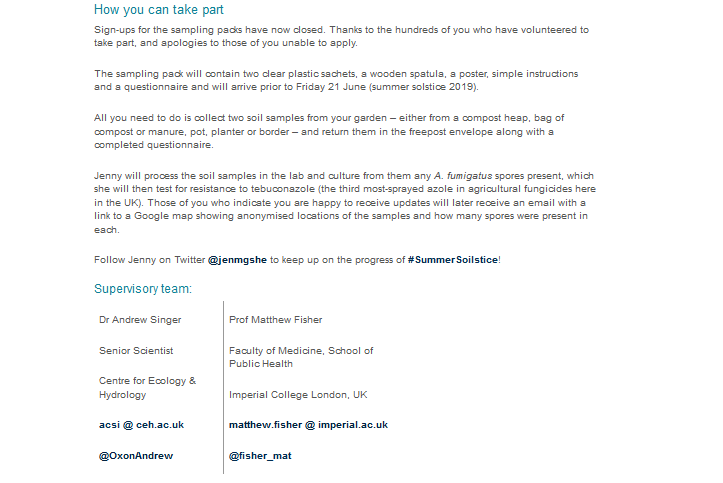

Supplementary Figure 8: Google form sent out four days after the first air sampling round asking participants for feedback on their sampling experience.

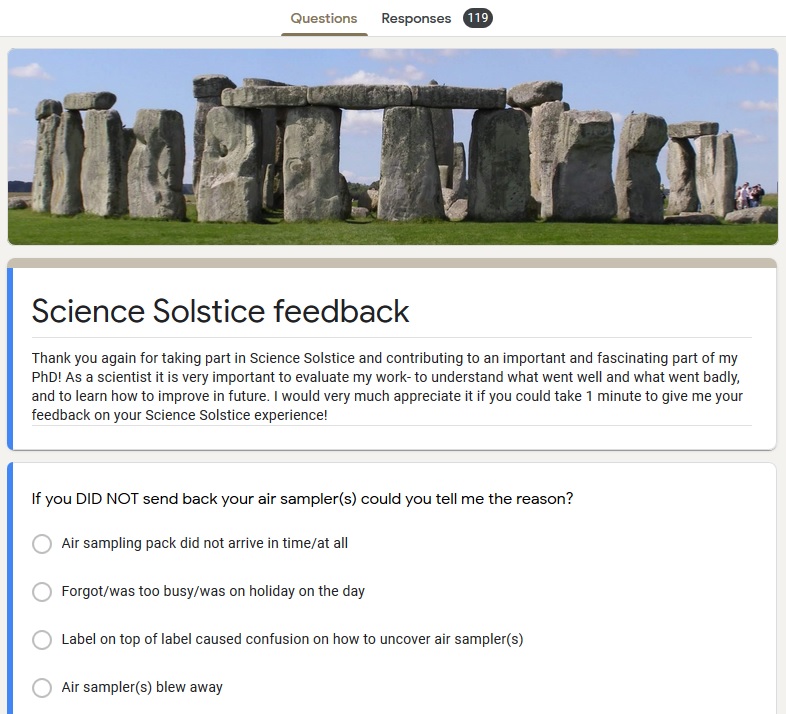

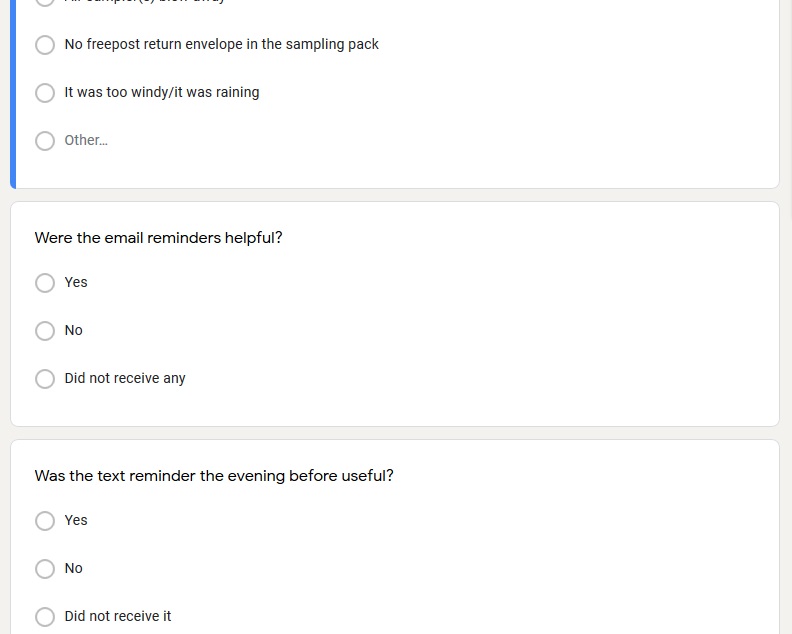

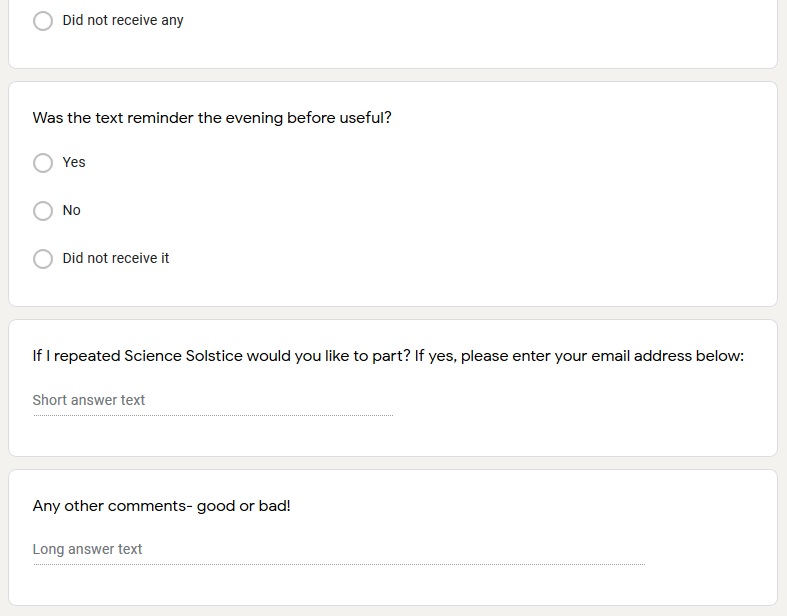

Supplementary Figure 9: Texts sent the day before the first and second air sampling dates to participants who had provided mobile numbers reminding them to take part in sampling the next day.

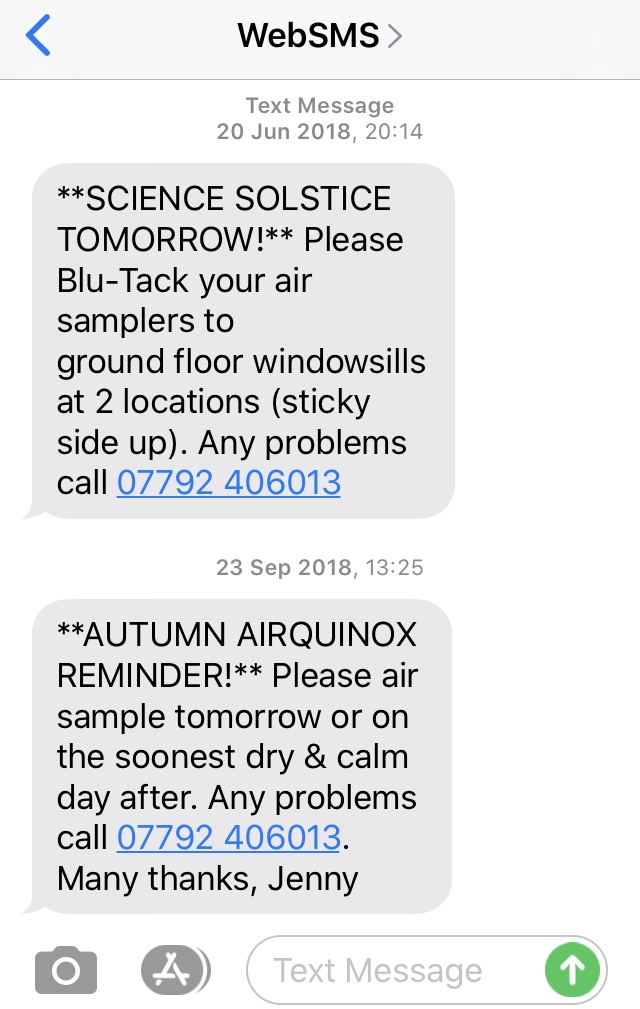

| Supplementary Figure 10: Timeline of changes made to air sampling methodology across the four air sampling rounds as a result of participant feedback. | | | |
| --- | --- | --- | --- |
| 1^st^ air sampling round:  21^st^ June 2018 | PROBLEM:  Several participants remove sticky labels stuck to the back of air samplers, instead of exposing the air sampler correctly. | PROBLEM: Participants report their air samplers blew away because adhesive putty was insufficiently adhesive. | PROBLEM:  Several participants contact author to apologise for not sampling because they were too busy on the sampling date. |
| 2^nd^ air sampling round:  24^th^ September 2018 | SOLUTION:  Location numbers handwritten onto back of air samplers instead of using sticky labels. | SOLUTION:  Double-sided foam pads were attached to the back of air samplers instead of adhesive putty. | SOLUTION:  For subsequent air sampling rounds participants were asked to sample on sampling date when possible, otherwise +/- 3 days. |
|  |  | PROBLEM: Participants report that foam pads leave residue on windowsills that requires chemical removal. |  |
| 3^rd^ air sampling round:  21^st^ December 2018  4^th^ air sampling round:  20^th^ March 2019 |  | SOLUTION:  Both adhesive putty and double-sided foam pads included in sampling packs for participants to choose which they use. |  |

Supplementary Table 1: Participant feedback provided in the additional comments box of the Google form sent out on 25^th^ June 2018.

1. More samplers as a contingency if/when we had problems with the first two
2. A practice experiment with some people to find out what could go wrong

After removing white covers I paper-clipped these face-to-face so as to keep uncontaminated until the evening when they were replaced on the sticky sheets. I do not know where other participants kept theirs (e.g. face-up or face-down / on windowsill or in a drawer) and it might not make much difference as long as they were inside the house (most likely, or they would have blown away!) but it might be something to consider?

All went OK. Suggest using thicker plastic as the hot weather tended to curl up the corners making it easier to blow away. It would be nice to have a dot map showing where the air samplers were located by volunteers etc. No other issues - you will sort out the 'label' problem so fingers crossed that you get the results you want. PS worked on Aspergillus fumigatus isolated from self-heating coalspoil sites 50 years ago so am interested in what you find anyway.

As spare sampler in the pack to replace any that went bad. Double sided tape as well as blue tac

Best wishes with your PhD project! I think it's excellent to use the general public to facilitate data collection. Well done!

blue tack didn't stick to UPVC very well

Difficult to leave the sticker unattended especially at the public area - Risk to be removed by cleaners etc

Enjoy the sampling.

Enjoyed taking part, thanks. I managed to get a finger print on one of my samples (sorry) - actually realised (too late) that there was a non sticky part of the backing paper from which we were presumably supposed to peel, but as this wasn't marked, I didn't realise this until after I'd put out the samples. Could this be marked somehow to make the unpeeling easier?
Also, wondering if these same samples could be used for other studies, e.g. looking at pollen counts/pollen species to develop/refine pollen maps for epidemiology research?
Anyway, good luck with the analysis, I'll look forward to hearing more in due course!

Everything was well explained.

Exciting project! We look forward to hearing about what you find!

Firstly, this is a great project, and I hope you get some nice results from it.

I don't know how feasible this is, but you might want to consider adding an edge around the sampler to hold on to, so people don't have to be concerned about the fingerprint and contamination issue.
From an ecological perspective, I suggest to include replicates for each location.

From what I could gather from your e-mails, I guess you will end up with "clumped data", i.e. mostly around Oxford, maybe some other UK science hotspots ;-), and then Leipzig and Ontario? It would be really interesting to see how you deal with this in the interpretation/analysation of the data.
And congrats on choosing a fascinating project, and a great citizen science approach to boot!

Fun to take part!

Fun! Next time I'll be back in Kent.

Good! Also I think I didn't sign up for updates. Please add me :)

Great project - was a tiny bit confused by the white backing at first, but your instruction sheet helped with that. Otherwise, the issue of blutack and making it stick to surfaces as I didn't want to touch the sticky top. Otherwise, very well communicated and explained what I needed to do.

Great project! Good luck. Hope our samples are useful (the cat stood on one of them!). Happy to help with anything in the future.

Great to be involved. We should be doing more of citizen science in general!

Great to take part.. Feel that I contaminated my sample through a level of ignorance over removal of label etc.. Next time make it ultra idiot proof.. You did to an extent but silly people on the school run may have struggled.. Sorry!

Happy to help. Good luck.

Home air sampler left out much longer than work air sampler - held up at work so air sampler at home was left out for just over the maximum time. I hope this won't affect / influence anything too much.
I thought the instructions were really clear, you provided a great amount of background information and made volunteers feel a part of the project.

I agree it was difficult not to get fingerprints on it

I also struggled with the fingerprint thing. Possibly having a marked edge that it was okay to touch could have helped with this? Also I emailed about not having downstairs windowsills to which you replied promptly and so I used other objects, this may have been outlined somewhere in the information but I didn't (and still haven't) read it properly. Just looked at the basic instructions...

I enjoyed taking part and look forward to seeing the results.

I had no problems removing the backing or with the sampler blowing away - even though it was a windy day.

I look forward to seeing your results. Could you let me know if my samples arrived with you please. Be great to see pics of what my samples look like on the plates. Would you like me to send you pics of the two sites to add to your photo collection. (Will also send this in an email in case you don't look at feedback for a while. )

I really like this experience and will definitively be happy to help in the future if needed!

I think I would maybe add a replacement sampling thingy, since then if the first one is screwed up, or one blew away, you could still have 2 samples from each place...

I think this was really well organised and you’ve done a brilliant job. But one thing I have learnt over the years, is to never underestimate people’s creativity in making mistakes or misunderstanding! You can be crystal clear and someone will always muck it up so don’t take it personally. All the best with the PHD.

I think you have been very diligent but I also think you have not captured as much associated data as you could have e.g. start and end time of sample while unlikely to have influenced results may have been useful as I expect some people did not stick to the 6-8 hours (I left home 8 am and did not return until 10 pm) , local weather (I would have provided tick box of possibilities with a fee text box comments), proximity to factors which might effect results e.g. arable agriculture fields with high probability of chemical sprays or what ever. Good luck with the PhD

I think you've had enough feedback on the tricky bits already. My samplers had a non-sticky edge (not sure if they all do), and it was easiest to peel from that edge.

I thought comms were good, and excellent excitement generated on Twitter!
I wonder if numbered steps *on* the collector might have helped avoid fingerprints (like mine!) - I.e. (printed on film). "1. Stick this side down to windowsill" or similar. (on white side) "2. peel this bit off and retain". As you would see on phone screen protectors.
I know it shouldn't be hard but I managed to do it the wrong way round, and I spent years in a micro lab...!
Well done for thinking up such an interesting experiment -looking forward to the results!

I thought it worked really well but the blu-tack is not suited for dusty or wet surfaces outside so I'd strongly suggest double-side foam tape cut into square (it has a peelable backing) it's what I used and it stayed just fine. I also found having a little note on the underside facing up saying 'Air-sampling please don't touch' helped.

I thought the instructions were very clear - even the white label! Was wondering why two samples and whether you needed these to be far apart or whether two very close together (e.g. as replicates) would be useful? Or having a control (inside) to compare with?

I thought the label thing was clear and obvious (particularly with the photos) but my friend, who did sample 2 for me in his flat, somehow found it very confusing. Noting to not do it overnight would have been helpful. The two sample thing may have put some people off as some people might not have two places to do it - I could not do one at work as I work in an upper floor office. I really enjoyed taking part though as a non-scientist - a simple task and I'm really interested in the results. I liked the photo instructions and the info sheet. The poster didn't seem that worthwhile though because by the time we received it it would have been a bit late for other people to take part.

I was happy to be a part of it and am looking forward to the results!

I was worried about 'effort' because I was able to put my sampler at home out very early in the morning and then got home super late, whereas at work I had meetings in the morning that meant it didn't get put out till 11 and taken in as I caught train home, so the time taken to collect the particles was mismatched with my work sample. For surveys I know 'effort' matters and it might be similar here. Can we have a bit to fill in how much time they were out or does that not matter? Otherwise, super easy to do and think it’s a great idea!

It was clearly explained and easy to do, except for the sticky labels, which were difficult to peel off.

It was tricky to remove the white backing without touching the sticky surface - I think including more Blu-tack would solve that.

it's hard to set air samples for 8 hours in two different locations; even if it's home and work it is unusual to have the exact time for both; perhaps more info on the allowance for time (like between 6 and 10?) would have helped

Just unfortunate that both samplers blew away :(

Look forward to seeing results!

Also, might be good to tell people to wipe the window ledge before sticking the thing. One of mine was so covered in pollen that it made the sticky surface immediately non sticky... the window sill needs to be pretty clean and dry.

Good luck!

more Blu-Tack :)

My outside windowsills are made of rough stone so blue tack doesn't work. A bit more information about why windowsills were suggested would have helped me choose an alternative spot. South Uist is Very Windy!
I struggled a bit to get the sticky surface completely re-covered after the sampling. Making the backing bigger than the sticky sampler might help.
Having said all that, it was easy enough and I would do it again.

Nice citizen science project!

nice work, well done; perhaps indeed try to find a solution for the blu Tack, not sticky enough (for eg stone)

no confusion about what label to remove, you’ve explained very well!

No issues. Was a doddle.

Nothing particular apart from that I felt the reminder text came in a bit late in the day, as I received the letter at my workplace, and forgot to bring it home the day before, so I had to return home on the Summer Solstice to deploy it. I think an e-mail/text reminder at noon on the day before and a text reminder in the evening is better. Good luck with your research!

Noticed about the finger prints and tried to avoid it by pressing down the Blu tack with the platic backing which seemed to work. Was fun, would like to see results in any way possible

Only comment would be something that you have already identified in your email, that it was quite difficult to avoid getting finger prints on the clear sticky screen

Perhaps next time make the peely away thing a different colour from the label so people aren't confused. Enjoyed doing the experiment and made me feel very important!

Please keep me on your email list for updates on project results.

Really enjoyed being part of it

Really enjoyed being part of your survey - looking forward to hearing about the findings. Got my granddaughter involved to provide the second site in town as I'm retired! Best wishes, Mary

Really enjoyed taking part - and reading the science bit on the information sheet. Great for my son to learn about too

Really fun and easy project, and your enthusiasm throughout was great!

Really nice and interesting experiments, easy to participate even far from your lab! I had just a question on the fact that it was impossible for me to not touch the sampler with my fingers. May it bias the results!?
Would be happy to follow your results.
Good luck.

Really well organised and sounds a great project - good luck with the analysis!

so wonderful to be a part of a fascinating project

sorry I think we contaminated the sample with fingerprints, I must have missed that in the instructions, other than that it was very well organised and simple to take part

The instructions were very clear and easy to understand. Really satisfying to be involved in some research! Best of luck with your studies and I look forward to seeing the results.

The three points in your email (confusion over labels etc) were all my experience too!

There is a large community of home educating families who love citizen science and use it for study. If you’re looking for even more people let me know and I can help to publicise it within our community groups. Additional info with experiments/sampling kits is great to help the children to learn. Good luck with your studies and career

Unfortunately I was one of the klutzs who peeled off the white address label rather than the whole top cover - pretty stupid, but instructions need to be foolproof, and it needs to be remembered that most of the people involved know nothing (as in NOTHING) about air sampling and hence aren't well equipped to know they have done something daft. Hope you got good coverage regardless - and get useful results.

Very impressed by your commitment to keeping people informed.

Very interesting project

Very interesting project and delighted to take part. As you are already aware, there was an issue about which label to peel off, though to be fair to you, instructions were provided in the pack; for next time, better to make it idiot-proof! All good wishes for your research.

Very interesting project. Great fun

We also struggled not to get finger prints on the sticky side but other than that everything was fine and the instructions were clear.

We love to get involved with physical hands on activities - so much more fun than simply filling in a survey. Well done and all the best with your research.

Well done, envelope with two samples on its way.

Were you still able to use the experiment even though some people took the wrong label off at first, but exposed the correct label after getting your later email?

What fun!

Will be interested to hear the results

Would be interested in finding out your results

Would definitely be interested in seeing the results (fellow scientist here...)

Yeah I think you mentioned before, but I got confused which location was which sample for a second, but luckily I'd filled out the info sheet before I stuck them out so I knew which was which. But if I hadn't very likely I would have forgotten.
